# 1,25-dihydroxyvitamin D3 potentiates the innate immune response of peripheral blood mononuclear cells from Japanese Black cattle

**DOI:** 10.1101/2023.04.29.538796

**Authors:** Youki Oyamada, Ei’ichi Iizasa, Amane Usa, Konosuke Otomaru

**Affiliations:** Division of Psychosomatic Internal Medicine, Department of Social and Behavioral Medicine, Graduate School of Medical and Dental Sciences, Kagoshima University, Japan; Joint faculty of veterinary medicine, Kagoshima University, Kagoshima, Japan

**Keywords:** Innate immunity, Japanese Black cattle, peripheral blood mononuclear cells, vitamin D

## Abstract

1,25-dihydroxyvitamin D_3_ (1,25(OH)_2_D_3_), a bioactive Vitamin D, is known to regulate immune responses in mammals. However, its impact on the innate immune responses of Japanese Black cattle, which are beef cattle endemic to Japan, remains unknown. Thus, in this study, we investigated the effect of 1,25(OH)_2_D_3_ on the immune responses of PBMC from Japanese Black cattle. PBMC were cultured with or without 1,25(OH)_2_D_3_ for measurement of cell viability, and stimulated with or without 1,25(OH)_2_D_3_ and lipopolysaccharide (LPS) for measurement of the gene expressions. As the results, the treatment of 1,25(OH)_2_D_3_ increased the cell viability. It also upregulated antibacterial peptides, *DEFB10* and *LAP* with or without LPS stimulation. Moreover, 1,25(OH)_2_D_3_ enhanced the inflammatory responses, *CXCL8* with LPS stimulation and *NOS2* with or without LPS stimulation, while reducing the expression of anti-inflammatory cytokine *IL10 w*ith or without LPS stimulation, leading to an inflammatory phenotype. However, in contrast to humans and mice, 1,25(OH)_2_D_3_ did not alter the expression of *TNF* and downregulated *TREM1* with LPS treatment. These results suggest that 1,25(OH)_2_D_3_ potentiates the innate immune responses of Japanese Black cattle, albeit with different effects and mechanisms as compared to humans and mice.

## 1. Introduction

The Innate immune response is elicited by the detection of pathogen-associated molecular patterns (PAMPs), which are derived from invading pathogens. In bovine peripheral blood mononuclear cells (PBMC), innate immune cells, such as monocytes, macrophages, and neutrophils, detect PAMPs through pattern recognition receptors (PRRs) including toll-like receptors (TLRs).

Lipopolysaccharide (LPS), which is one of PAMPs derived from gram-negative bacteria, is detected by TLR4 to trigger various innate immune responses, such as producing antibacterial peptides, inflammatory cytokines, and bactericidal nitric oxide (NO) (Hazlett & Wu, 2011).

Vitamin D is a fat-soluble vitamin for the formation of bones and teeth and for the absorption of phosphorus and calcium in the intestines (Eder & Grundmann, 2022). Vitamin D_3_ is produced from its precursor 7-dehydrocholesterol in the skin by sunlight (UV-B) or food intake. vitamin D_3_ is further transformed into its bioactive metabolite, 1,25-dihydroxyvitamin D_3_ (1,25(OH)_2_D_3_) in liver and kidneys. In recent decades, many reports have also highlighted the critical role of vitamin D in regulating the innate immune responses in humans and mice (Almeida Moreira Leal et al., 2020; Colotta et al., 2017; Huang & Huang, 2021; Khoo et al., 2011). Most reports have shown 1,25(OH)_2_D_3_ treatments reduce inflammatory cytokine production in humans and mice. For instance, 1,25(OH)_2_D_3_ inhibits LPS-induced IL-6 and TNF production in human monocytes and mouse bone marrow-derived macrophages via activation of MAPK phosphatase-1 (MKP-1) (Zhang et al., 2012). 1,25(OH)_2_D_3_ also prevents NF-κB activation for upregulation of Cox-2 and inflammatory cytokines in RAW264.7, a murine macrophage cell line, when stimulated by LPS (Chen et al., 2013; Q. Wang et al., 2014). In contrast, some reports have shown the 1,25(OH)_2_D_3_ enhances the LPS-induced inflammatory cytokines in human and murine cells (Di Rosa et al., 2012; Kim et al., 2013; Rigo et al., 2012). In addition, antibacterial peptides and iNOS, which produces NO, are upregulated by 1,25(OH)_2_D_3_ in human cells both in the presence and absence of LPS (Nelson et al., 2012). 1,25(OH)_2_D_3_ also confers bactericidal activity against mycobacteria by inducing the expression of antibacterial peptides, such as a cathelicidin and β-defensins (BDNFs) (Nelson et al., 2012).

Furthermore, 1,25(OH)_2_D_3_ induces the expression of CD14 and TREM1, which are the molecules enhancing TLR4 signaling (Hosoda et al., 2015; Kim et al., 2013; Koivisto et al., 2020; Nelson et al., 2012; Rigo et al., 2012). Therefore, the induction of these genes may account for the upregulation of LPS-induced immune responses by 1,25(OH)_2_D_3._

On the other hand, despite limited investigations into the impact of 1,25(OH)_2_D_3_ on bovine immunity, the previous reports suggest that 1,25(OH)_2_D_3_ could enhance innate immune responses in cattle. Specifically, 1,25(OH)_2_D_3_ upregulates iNOS and RANTES genes but not any bovine cathelicidin genes in the monocyte from Holstein dairy cattle (Nelson et al., 2010). Additionally, RNA-seq and sequential quantitative real-time PCR (qRT-PCR) have also revealed that 1,25(OH)_2_D_3_ upregulates bovine BDNFs, while not affecting lingual and tracheal antimicrobial peptides (LAP and TAP, respectively) in Holstein dairy cattle (Merriman et al., 2015).

On the other hand, the impact of 1,25(OH)_2_D_3_ on immune responses in beef cattle remains largely unexplored. Given that dairy and beef cattle have been selectively bred for distinct traits, namely milk and meat quality, respectively, the difference in their genetic traits and farming practices have possibly impacted immunity. Indeed, some studies have reported the difference between dairy and beef calves in immune responses (Johnston et al., 2020; Surlis et al., 2018).

More than 90% of the beef cattle raised in Japan are Japanese Black cattle. Japanese black cattle are a breed of beef cattle, which have specific traits and are endemic to Japan (Motoyama et al., 2016). Therefore, it is crucial to investigate the effect of vitamin D on the immune response of Japanese Black cattle to compare it to dairy cattle and other mammals. In this study, we purified the PBMC of Japanese Black cattle and examined the effect of vitamin D on survival *in vitro* culture. Additionally, we treated PBMC with 1,25(OH)_2_D_3_ in the absence and presence of LPS to examine the effect of vitamin D on the innate immune response of PBMC.

## 2. Materials and methods

### 2.1 Animals

All Japanese Black cattle used in this study were reared on farms located in Kagoshima Prefecture, Japan. Following their birth, the calves were permitted to remain with their dams for one week before being segregated into individual hutches until they reached two weeks of age. From three weeks to twelve weeks of age, the calves were grouped and housed in paddocks consisting of 18 individuals each and provided with a milk replacer as a dietary supplement. Upon reaching 13 weeks of age, the calves were weaned and placed in the paddocks of six individuals. The calves received equivalent treatment and nutrition to satisfy the nutritional criteria of the Japanese beef cattle feeding standard (MAFF 2008). All experimental procedures were reviewed and approved by the Kagoshima University Laboratory Animal Committee, Japan (Approval number: KVM220004).

### 2.2 PBMC collection

Blood was collected from the Jugular vein of the calves from 24-32 weeks of age using vacutainer heparin tubes. PBMC were purified as previously described (Maeda et al., 2011). Briefly, 10 mL of heparinized blood was diluted twofold in phosphate-buffered saline (PBS), placed above the separation medium solution (Lymphocyte Separation Medium 1077, Immuno-Biological Laboratories, Japan; specific gravity 1.077), and then centrifuged at 400 × g for 60 min at room temperature. Mononuclear cells at the interface were collected, washed twice with PBS by centrifugation at 400 × g for 5 min, and subsequently treated with 0.83% ammonium chloride. The cells were gently agitated for 5 min to remove any residual red blood cells. The isolated PBMCs were washed with PBS and suspended in RPMI1640 medium (Invitrogen, Tokyo, Japan) containing 100-U/mL penicillin G (Meiji Seika, Tokyo, Japan).

### 2.3 Cell viability in culture

Examination of cell viability in culture was conducted following previous reports (De Abreu Costa et al., 2017; Yang et al., 1995). The isolated PBMCs (n = 26) were dispensed into 48-well plates (Greiner Bio-One, Tokyo, Japan) at a density of 5 × 10^5^ cells / well, with a final volume of 1 mL / well in RPMI 1640 medium supplemented with 10% heat-inactivated fetal calf serum (FCS; Cansera International Inc., Rexdale, Canada). The cells were treated with 1,25(OH)_2_D_3_ (Calcitriol, Fujifilm Wako, Japan) at 37°C for 24 and 72 h in a humidified 5% CO_2_ atmosphere. The concentration of 1,25(OH)_2_D_3_ was set to 10^−8^ mol/L with reference to the previous reports (Villaggio et al., 2012). After the treatments, live cells were counted with trypan blue (0.5% -trypan blue stain, Nakai Tesque, Japan), and the cell viability was calculated by dividing the number of cells after 24 and 72 h treatment by that of 0 h.

### 2.4 *In vitro* PBMC culture and stimulation

The isolated PBMC (n = 21) were seeded at a density of 5 × 10^6^ cells/well in 48-well plates containing 1 ml/well of RPMI 1640 medium supplemented with 10% heat-inactivated FCS. The cells were cultured at 37°C in a humidified 5% CO_2_ atmosphere. To investigate the effect of 1,25(OH)_2_D_3_ on LPS-stimulated PBMC, the cells were stimulated by 1 μg/ml LPS in the presence or absence of 10^−8^ mol/L 1,25(OH)_2_D_3_ as previously reported (Villaggio et al., 2012). After 24 h of stimulation, the cells were harvested and suspended in Trizol (Thermo Fisher Scientific) for RNA extraction.

### 2.5 Quantative real-time PCR

Total RNA was extracted from PBMC (n = 21) using Trizol according to the manufacturer’s instructions. The total RNA was reverse-transcribed by ReverTra Ace qPCR RT Master Mix with gDNA Remover (TOYOBO, Japan). Quantitative real-time PCR (qRT-PCR) was performed using THUNDERBIRD NEXT SYBR qPCR Mix (TOYOBO) and StepOnePlus Real-Time PCR system. The primer sets (Merriman et al., 2015; Sassu et al., 2020; Takanashi et al., 2013; Tomasinsig et al., 2010; Yamane et al., 2008) used in the analysis are listed in Table 1. The relative gene expression was calculated by the ΔΔCt method, with GAPDH serving as an internal control.

**Table 1.**
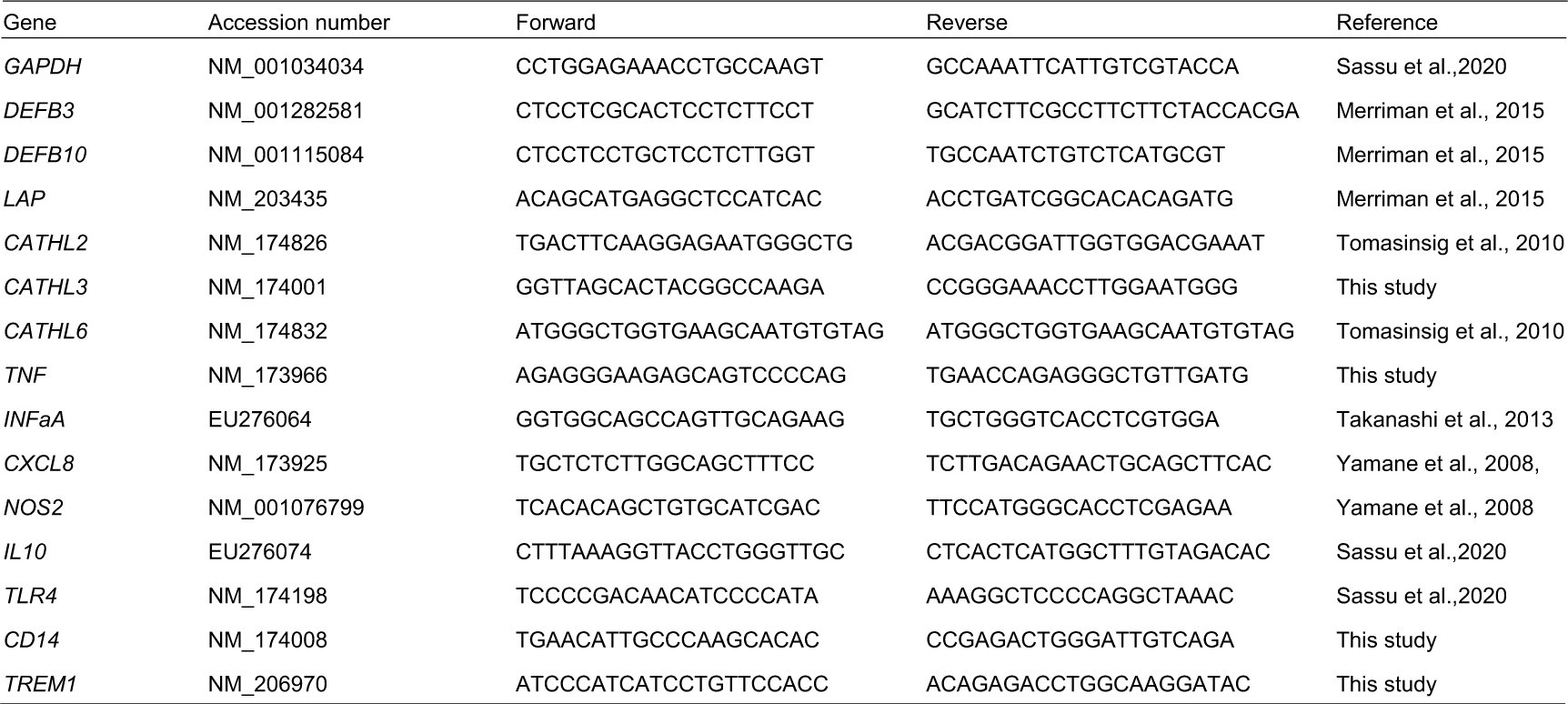
The primer sets used in this study

### 2.6 Statistical analysis

Statistical analyses were conducted using GraphPad Prism 9 software. Outliers were detected using the ROUT test and subsequently removed. Comparisons between two groups were made using the two-tailed unpaired *t*-test. Multiple groups were compared using one-way ANOVA with the Dunnett’s T3 post-hoc test. *P* value <0.05 was considered statistically significant.

## 3. Results and discussion

Prior to investigating the influence of vitamin D on immune responses, we verified that on the viability of bovine PBMC. The cells were isolated from Japanese Black calves and then cultured with or without the bioactive vitamin D, 1,25(OH)_2_D_3_. The cells were counted, and the viability was calculated at 24 and 72 h. The result indicates a significant difference in cell viability with the presence of 1,25(OH)_2_D_3_ at 72 h but not at 24 h (Fig. 1). This result suggests that vitamin D can increased the viability of PBMC. Although the precise mechanism and effect of vitamin D on cell viability remain unclear, a previous study has reported that vitamin D treatment inhibits the apoptosis of PBMC from human SLE patients by upregulation of an antiapoptotic molecule *Bcl-2* (Tabasi et al., 2015). Hence, we speculate that a similar phenomenon could occur in the bovine PBMC. Moreover, since 1,25(OH)_2_D_3_ exhibited no cytotoxicity on PBMCs after 24 h treatment (Fig. 1). Thus, we subsequently evaluated the effect of 1,25(OH)_2_D_3_ on immune responses at 24 h in the following experiments.

**Figure 1.**
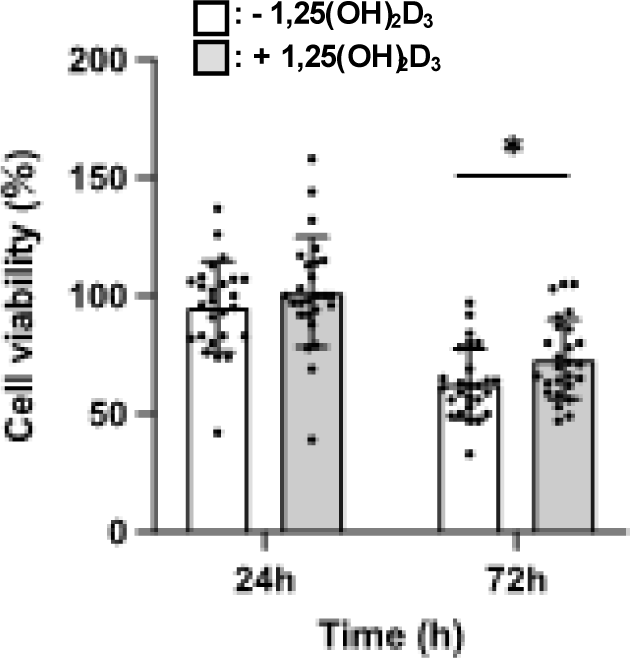
The effect of 1,25(OH)_2_D_3_ on the viability of bovine PBMC. Bovine PBMC were treated 1,25(OH)_2_D_3_ (10^−8^ ng/mL) for 24 and 72 h. Cells were counted and the cell viability was calculated. Data are presented as means ± SEM (n = 26) and Statistical comparison was performed using the Student *t-*test or one-way ANOVA with t the Dunnett’s T3 post hoc test. \**p* < 0.05.

Previous studies have demonstrated that vitamin D treatment of human and Holstein cattle cells leads to an elevation of β-defensin and other antimicrobial peptide productions (Kościuczuk et al., 2014; Merriman et al., 2015; T.-T. Wang et al., 2004). Thus, we investigated the impact of 1,25(OH)_2_D_3_ on the expression of these genes in PBMC from Japanese Black cattle. Following treatment of PBMC with 1,25(OH)_2_D_3_ in the presence or absence of LPS, qRT-PCR quantified the expression of β-defensins (DEFBs), such as *DEFB3*, *DEFB10*, and *LAP*, and cathelicidins, such as *CATHL2* (*Bac5*), *CATHL3*, and *CATHL6*. The results showed that 1,25(OH)_2_D_3_ upregulated the expression of *DEFB10* and *LAP* both in the presence and absence of LPS stimulation, while having no effect on that of *DEFB3* (Fig. 2). Furthermore, our findings indicated that it treatments did not affect the expression of any cathelicidins (data not shown), consistent with a previous study of monocytes in Holstein cattle (Nelson et al., 2010). Previous studies have shown that the expressions of a cathelicidin antimicrobial peptide (CAMP) was upregulated by 1,25(OH)_2_D_3_ in human (Liu et al., 2006; T.-T. Wang et al., 2004). Thus, our findings highlight the differences between humans and cattle. CAMP, also known as LL-37, is the only member of the cathelicidin family protein present in human genome and its promoter contains the vitamin D receptor binding element (VDRE) (Dürr et al., 2006). However, the VDRE in the CAMP gene originated in the primates including human and is absent in other mammals (Gombart et al., 2009). Moreover, it is known that there are about 10 cathelicidin genes in cattle that have evolved distantly from those in humans (Laroche et al., 2013). Therefore, our results can be consistent with this evolutionary divergence. On the other hand, a previous study reported that 1,25(OH)_2_D_3_ increased the expression of *DEFB3* and *DEFB10* but not that of *LAP* in Holstein cattle monocytes. Thus, these results also demonstrate that there is also a subtle difference between Japanese Black cattle and Holstein cattle in the impact of 1,25(OH)_2_D_3_.

**Figure 2.**
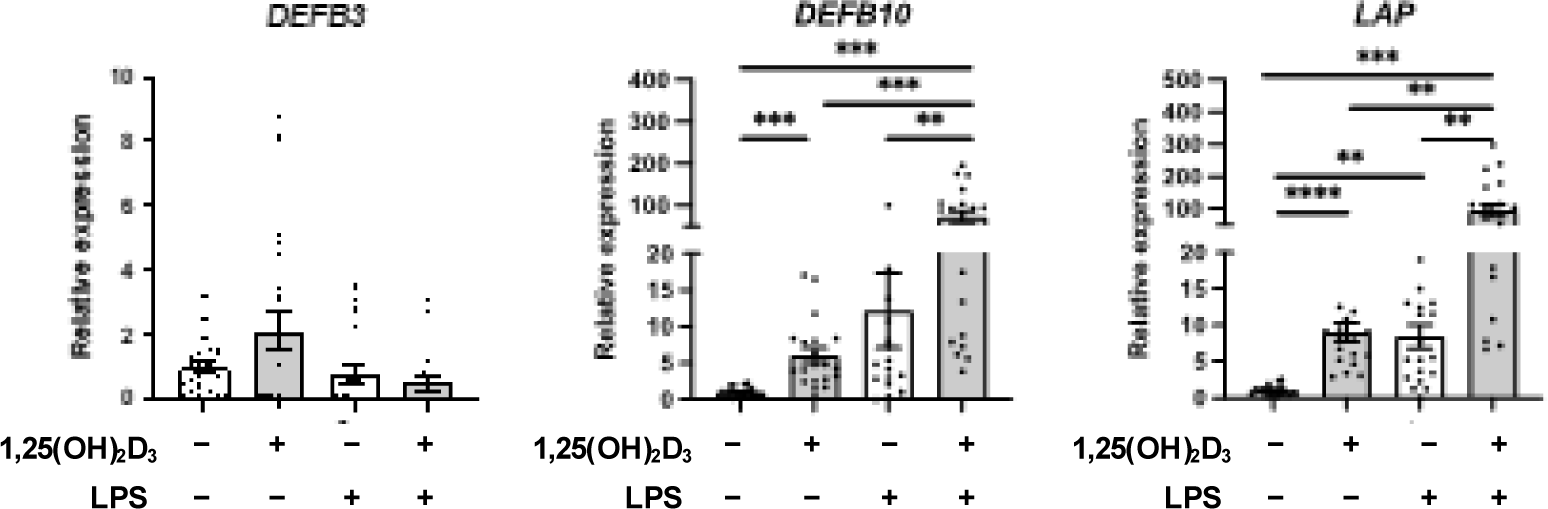
1,25(OH)_2_D_3_ treatment enhances the expression of antibacterial peptides. PBMC were stimulated by LPS (1 μg/ml) with or without 1,25(OH)_2_D_3_ (10^−8^ ng/ml) for 24 h. Relative expression of the DEFBs (*BNBD3*, and *BNBD10*) and *LAP* was determined by qRT-PCR. Data are presented as means ± SEM (n = 17-21) and Statistical comparison was performed using one-way ANOVA with the Dunnett’s T3 post hoc test. \*\*\*\**p* < 0.0001, \*\*\**p* < 0.001, \*\**p* < 0.01.

We next examined the impact of 1,25(OH)_2_D_3_ on the expression of cytokines and *NOS2* in PBMC under LPS stimulation. The results indicate that 1,25(OH)_2_D_3_ upregulated the LPS-induced expression of *CXCL8* (*IL8*) and *NOS2*, while it downregulated that of *IL10* (Fig. 3). In contrast to previous studies in humans and mice (Khoo et al., 2011; Zhang et al., 2012), 1,25(OH)_2_D_3_ did not affect that of *TNF* in the bovine PBMC (Fig. 3). CXCL8 is a chemokine that recruits neutrophils and plays a vital role in the eradication of bacterial infections (Cambier et al., 2023). Meanwhile, the *NOS2* gene encodes iNOS, which generates NO that is crucial for eliminating intracellular bacteria such as mycobacteria (Boylan et al., 2020). Thus, these results show that 1,25(OH)_2_D_3_ potentiates an inflammatory phenotype that promotes the clearance of bacteria. Conversely, 1,25(OH)_2_D_3_ had no effect on the expression of type I Interferon *IFNaA*, which restricts viral replication (McNab et al., 2015), suggesting that 1,25(OH)_2_D_3_ is less effective for viral infections. In Holstein monocytes, the expression of *NOS2* and *RANTES* is reported to be also enhanced by vitamin D in response to PAMPs (Nelson et al., 2010, 2011). Accordingly, we hypothesize that 1,25(OH)_2_D_3_ may specifically regulate the expression of *NOS2* and chemokines, including *CXCL8* and *RANTES*, in cattle.

**Figure 3.**
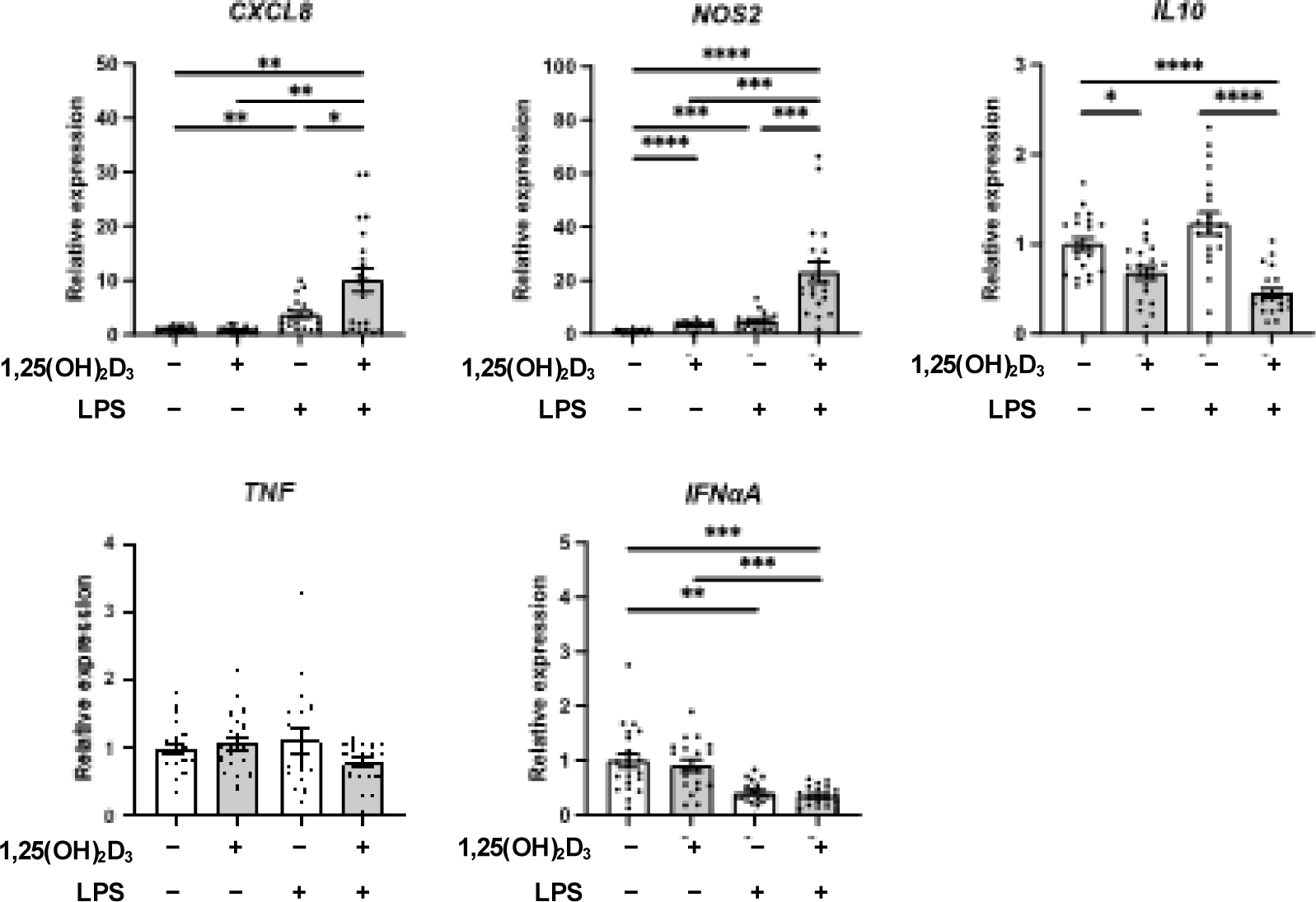
1,25(OH)_2_D_3_ upregulates the expression of *CXCL8* and *NOS2* while downregulates that of *IL10*. PBMC were stimulated by LPS (1 μg/ml) with or without 1,25(OH)_2_D_3_ (10^−8^ ng/mL) for 24 h. Relative expression of *CXCL8*, *NOS2*, *IL10*, *TNF*, and *IFNαA* was determined by qRT-PCR. Data are presented as means ± SEM (n = 18-21) and Statistical comparison was performed using one-way ANOVA with the Dunnett’s T3 post hoc test. \*\*\*\**p* < 0.0001, \*\*\**p* < 0.001, \*\**p* < 0.01, \**p* < 0.05.

Previous reports have shown that 1,25(OH)_2_D_3_ enhances the expression of *CD14*, which facilitates LPS transfer to TLR4, in human monocytes (Schauber et al., 2007; Scherberich et al., 2005). Although the mechanisms underlying the immune regulation by vitamin D remain unclear, it is considered one of the possible mechanisms for the upregulation of inflammatory genes. Therefore, to investigate the mechanism in cattle, we evaluated the expression of *TLR4* and *CD14* in PBMC from Japanese Black cattle treated with 1,25(OH)_2_D_3_ in the presence or absence of LPS. While LPS stimulation significantly decreased the expression of these genes, 1,25(OH)_2_D_3_ affected the expression of neither *TLR4* nor *CD14* in bovine PBMC (Fig. 4A). Another possible mechanism involves the upregulation of TREM1 expression by 1,25(OH)_2_D_3_. *TREM1* is one of Triggering Receptor Expressed on Myeloid cells (TREM) family receptors that are known to amplify LPS-induced and other inflammations (Bouchon et al., 2001; Tammaro et al., 2017). Although its physiological ligands remain unknown, the cross-linking of the receptor by antibodies and simultaneous activation of PRRs leads to the upregulation of immune responses (Tammaro et al., 2017). Previous reports have shown that 1,25(OH)_2_D_3_ induces the expression of TREM1, and followed by its cross-linking and LPS stimulation enhances cytokine expressions in human cells (Carrasco et al., 2019; Hosoda et al., 2015; Kim et al., 2013; Rigo et al., 2012).

**Figure 4.**
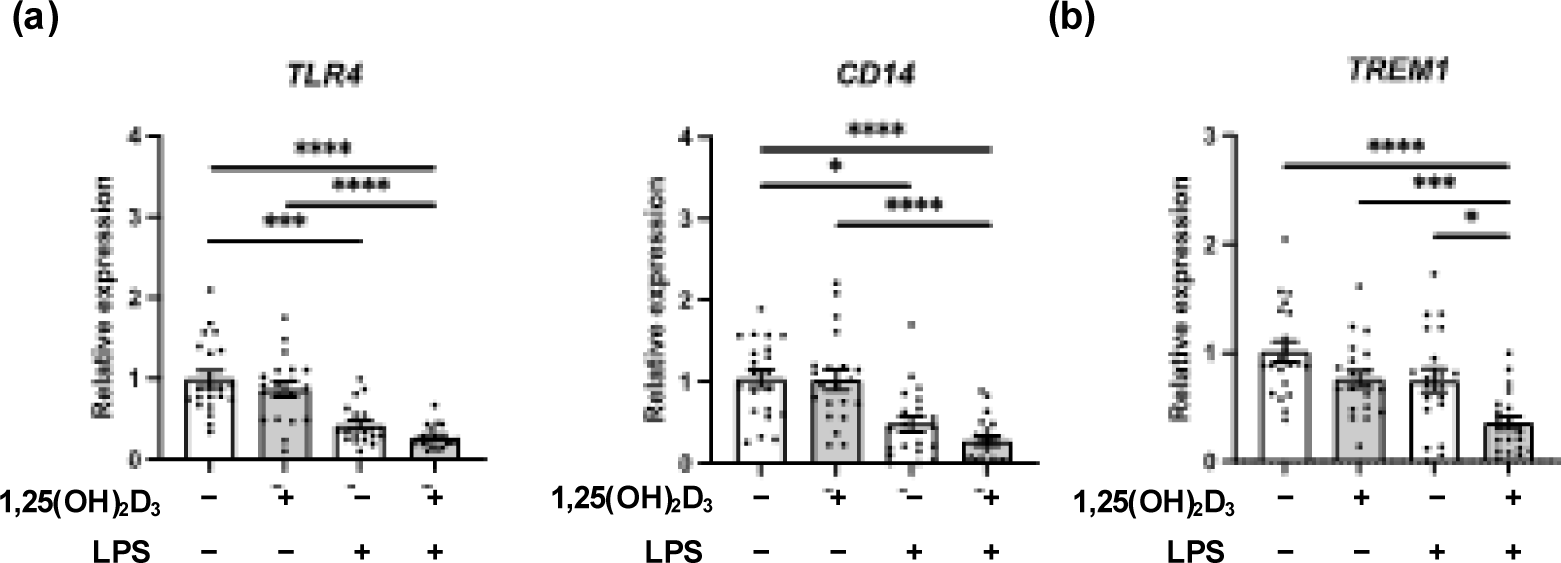
1,25(OH)_2_D_3_ treatment did not affect the expression of *TLR4* and *CD14* but downregulated that of *TREM1*. (a, b) PBMC were stimulated by LPS (1 μg/ml) with or without Vitamin D (10^−8^ ng/ml) for 24 h. Relative expression of the *TLR4*, *CD14* (a), and TREM1(b) was determined by qRT-PCR. Data are presented as means ± SEM (n = 19-21) and Statistical comparison was performed using one-way ANOVA with Dunnett’s T3 post hoc test. \*\*\*\**p* < 0.0001, \*\*\**p* < 0.001, \**p* < 0.05.

Therefore, we next assessed the expression of *TREM1* in bovine PBMC following 1,25(OH)_2_D_3_ treatment in the presence or absence of LPS. In contrast to humans, 1,25(OH)_2_D_3_ treatment of bovine PBMC with LPS significantly decreased the expression of *TREM1* (Fig. 4b). This is the first report investigating 1,25(OH)_2_D_3_ effect on the expression of *CD14* and *TREM1* in cattle. These results suggest that 1,25(OH)_2_D_3_ treatment potentiates the inflammatory response of cattle through a mechanism different from that in humans. On the other hand, we cannot exclude the possibility that the time course of the analysis for the expression of these genes was not appropriate. As the expression of *TLR4* and *CD14* is reduced at 24 h after treatment (Fig. 4a), it is possible that the gene expression of receptors and membrane proteins required for the early phase of the immune response is already reduced at this time point. However, the induction of *TREM1* expression in humans has been detected at similar times (24 h after treatments) (Hosoda et al., 2015; Kim et al., 2013; Rigo et al., 2012). Thus, at least, it suggests that there may be a different regulatory mechanism as compared to humans.

Overall, our study demonstrates that 1,25(OH)_2_D_3_ treatment increased the viability of and upregulates the expression of inflammatory genes of PBMC derived from Japanese Black cattle. Specifically, our findings reveal that *DEFB*s, *NOS2*, and *CXCL8* genes, which play pivotal roles in immune responses against bacteria, were significantly upregulated upon 1,25(OH)_2_D_3_ treatment. Our results are largely consistent with previous reports investigating the impact of 1,25(OH)_2_D_3_ on the immune responses in Holstein dairy cattle, indicating that its effects are comparable in beef and dairy cattle. Our finding suggests that vitamin D plays a pivotal role in potentiating the immune response in Japanese Black cattle.

## Acknowledgement

This study was supported by public interest incorporated foundation the Ito Foundation

## References

Almeida Moreira Leal, L. K., Lima, L. A., Alexandre de Aquino, P. E., Costa de Sousa, J. A., Jataí Gadelha, C. V., Felício Calou, I. B., Pereira Lopes, M. J., Viana Lima, F. A., Tavares Neves, K. R., Matos de Andrade, G., & Socorro de Barros Viana, G. (2020). Vitamin D (VD3) antioxidative and anti-inflammatory activities: Peripheral and central effects. European Journal of Pharmacology, 879, 173099. https://doi.org/10.1016/j.ejphar.2020.173099

Bouchon, A., Facchetti, F., Weigand, M. A., & Colonna, M. (2001). TREM-1 amplifies inflammation and is a crucial mediator of septic shock. Nature, 410(6832), Article 6832. https://doi.org/10.1038/35074114

Boylan, M., O’Brien, M. B., Beynon, C., & Meade, K. G. (2020). 1,25(OH)D vitamin D promotes NOS2 expression in response to bacterial and viral PAMPs in primary bovine salivary gland fibroblasts. Veterinary Research Communications, 44(2), 83–88. https://doi.org/10.1007/s11259-020-09775-y

Cambier, S., Gouwy, M., & Proost, P. (2023). The chemokines CXCL8 and CXCL12: Molecular and functional properties, role in disease and efforts towards pharmacological intervention. Cellular & Molecular Immunology, 20(3), Article 3. https://doi.org/10.1038/s41423-023-00974-6

Carrasco, K., Boufenzer, A., Jolly, L., Le Cordier, H., Wang, G., Heck, A. J., Cerwenka, A., Vinolo, E., Nazabal, A., Kriznik, A., Launay, P., Gibot, S., & Derive, M. (2019). TREM-1 multimerization is essential for its activation on monocytes and neutrophils. Cellular and Molecular Immunology, 16(5), 460–472. https://doi.org/10.1038/s41423-018-0003-5

Chen, Y., Liu, W., Sun, T., Huang, Y., Wang, Y., Deb, D. K., Yoon, D., Kong, J., Thadhani, R., & Li, Y. C. (2013). 1,25-Dihydroxyvitamin D Promotes Negative Feedback Regulation of TLR Signaling via Targeting MicroRNA-155–SOCS1 in Macrophages. The Journal of Immunology, 190(7), 3687–3695. https://doi.org/10.4049/jimmunol.1203273

Colotta, F., Jansson, B., & Bonelli, F. (2017). Modulation of inflammatory and immune responses by vitamin D. Journal of Autoimmunity, 85, 78–97. https://doi.org/10.1016/j.jaut.2017.07.007

De Abreu Costa, L., Henrique Fernandes Ottoni, M., Dos Santos, M. G., Meireles, A. B., Gomes de Almeida, V., De Fátima Pereira, W., Alves de Avelar-Freitas, B., & Eustáquio Alvim Brito-Melo, G. (2017). Dimethyl Sulfoxide (DMSO) Decreases Cell Proliferation and TNF-α, IFN-γ, and IL-2 Cytokines Production in Cultures of Peripheral Blood Lymphocytes. Molecules, 22(11), Article 11. https://doi.org/10.3390/molecules22111789

Di Rosa, M., Malaguarnera, G., De Gregorio, C., Palumbo, M., Nunnari, G., & Malaguarnera, L. (2012). Immuno-modulatory effects of vitamin D3 in human monocyte and macrophages. Cellular Immunology, 280(1), 36–43. https://doi.org/10.1016/j.cellimm.2012.10.009

Dürr, U. H. N., Sudheendra, U. S., & Ramamoorthy, A. (2006). LL-37, the only human member of the cathelicidin family of antimicrobial peptides. Biochimica et Biophysica Acta (BBA) - Biomembranes, 1758(9), 1408–1425. https://doi.org/10.1016/j.bbamem.2006.03.030

Eder, K., & Grundmann, S. M. (2022). Vitamin D in dairy cows: Metabolism, status and functions in the immune system. Archives of Animal Nutrition, 76(1), 1–33. https://doi.org/10.1080/1745039X.2021.2017747

Gombart, A. F., Saito, T., & Koeffler, H. P. (2009). Exaptation of an ancient Alu short interspersed element provides a highly conserved vitamin D-mediated innate immune response in humans and primates. BMC Genomics, 10(1), 321. https://doi.org/10.1186/1471-2164-10-321

Hazlett, L., & Wu, M. (2011). Defensins in innate immunity. Cell and Tissue Research, 343(1), 175–188. https://doi.org/10.1007/s00441-010-1022-4

Hosoda, H., Tamura, H., & Nagaoka, I. (2015). Evaluation of the lipopolysaccharide-induced transcription of the human TREM-1 gene in vitamin D3-matured THP-1 macrophage-like cells. International Journal of Molecular Medicine, 36(5), 1300–1310. https://doi.org/10.3892/ijmm.2015.2349

Huang, F.-C., & Huang, S.-C. (2021). Active vitamin D3 attenuates the severity of Salmonella colitis in mice by orchestrating innate immunity. Immunity, Inflammation and Disease, 9(2), 481–491. https://doi.org/10.1002/iid3.408

Johnston, D., Mukiibi, R., Waters, S. M., McGee, M., Surlis, C., McClure, J. C., McClure, M. C., Todd, C. G., & Earley, B. (2020). Genome wide association study of passive immunity and disease traits in beef-suckler and dairy calves on Irish farms. Scientific Reports, 10(1), Article 1. https://doi.org/10.1038/s41598-020-75870-4

Khoo, A.-L., Chai, L. Y. A., Koenen, H. J. P. M., Kullberg, B.-J., Joosten, I., van der Ven, A. J. A. M., & Netea, M. G. (2011). 1,25-dihydroxyvitamin D3 Modulates Cytokine Production Induced by Candida albicans: Impact of Seasonal Variation of Immune Responses. The Journal of Infectious Diseases, 203(1), 122–130. https://doi.org/10.1093/infdis/jiq008

Kim, T.-H., Lee, B., Kwon, E., Choi, S. J., Lee, Y. H., Song, G. G., Sohn, J., & Ji, J. D. (2013). Regulation of TREM-1 expression by 1,25-dihydroxyvitamin D3 in human monocytes/macrophages. Immunology Letters, 154(1), 80–85. https://doi.org/10.1016/j.imlet.2013.08.012

Koivisto, O., Hanel, A., & Carlberg, C. (2020). Key Vitamin D Target Genes with Functions in the Immune System. Nutrients, 12(4), Article 4. https://doi.org/10.3390/nu12041140

Kościuczuk, E. M., Lisowski, P., Jarczak, J., Krzyżewski, J., Zwierzchowski, L., & Bagnicka, E. (2014). Expression patterns of β-defensin and cathelicidin genes in parenchyma of bovine mammary gland infected with coagulase-positive or coagulase-negative Staphylococci. BMC Veterinary Research, 10(1), 246. https://doi.org/10.1186/s12917-014-0246-z

Laroche, F. J. F., Tulotta, C., Lamers, G. E. M., Meijer, A. H., Yang, P., Verbeek, F. J., Blaise, M., Stougaard, J., & Spaink, H. P. (2013). The embryonic expression patterns of zebrafish genes encoding LysM-domains. Gene Expression Patterns, 13(7), 212–224. https://doi.org/10.1016/j.gep.2013.02.007

Liu, P. T., Stenger, S., Li, H., Wenzel, L., Tan, B. H., Krutzik, S. R., Ochoa, M. T., Schauber, J., Wu, K., Meinken, C., Kamen, D. L., Wagner, M., Bals, R., Steinmeyer, A., Zügel, U., Gallo, R. L., Eisenberg, D., Hewison, M., Hollis, B. W., … Modlin, R. L. (2006). Toll-Like Receptor Triggering of a Vitamin D-Mediated Human Antimicrobial Response. Science, 311(5768), 1770–1773. https://doi.org/10.1126/science.1123933

Maeda, Y., Tanaka, R., Ohtsuka, H., Matsuda, K., Tanabe, T., & Oikawa, M. (2011). Comparison of the Immunosuppressive Effects of Dexamethasone, Flunixin Meglumine and Meloxicam on the *In Vitro* Response of Calf Peripheral Blood Mononuclear Cells. Journal of Veterinary Medical Science, 73(7), 957–960. https://doi.org/10.1292/jvms.10-0422

McNab, F., Mayer-Barber, K., Sher, A., Wack, A., & O’Garra, A. (2015). Type I interferons in infectious disease. Nature Reviews Immunology, 15(2), Article 2. https://doi.org/10.1038/nri3787

Merriman, K. E., Kweh, M. F., Powell, J. L., Lippolis, J. D., & Nelson, C. D. (2015). Multiple β-defensin genes are upregulated by the vitamin D pathway in cattle. The Journal of Steroid Biochemistry and Molecular Biology, 154, 120–129. https://doi.org/10.1016/j.jsbmb.2015.08.002

Motoyama, M., Sasaki, K., & Watanabe, A. (2016). Wagyu and the factors contributing to its beef quality: A Japanese industry overview. Meat Science, 120, 10–18. https://doi.org/10.1016/j.meatsci.2016.04.026

Nelson, C. D., Nonnecke, B. J., Reinhardt, T. A., Waters, W. R., Beitz, D. C., & Lippolis, J. D. (2011). Regulation of Mycobacterium-Specific Mononuclear Cell Responses by 25-Hydroxyvitamin D3. PLOS ONE, 6(6), e21674. https://doi.org/10.1371/journal.pone.0021674

Nelson, C. D., Reinhardt, T. A., Lippolis, J. D., Sacco, R. E., & Nonnecke, B. J. (2012). Vitamin D Signaling in the Bovine Immune System: A Model for Understanding Human Vitamin D Requirements. Nutrients, 4(3), Article 3. https://doi.org/10.3390/nu4030181

Nelson, C. D., Reinhardt, T. A., Thacker, T. C., Beitz, D. C., & Lippolis, J. D. (2010). Modulation of the bovine innate immune response by production of 1α,25-dihydroxyvitamin D3 in bovine monocytes. Journal of Dairy Science, 93(3), 1041–1049. https://doi.org/10.3168/jds.2009-2663

Rigo, I., McMahon, L., Dhawan, P., Christakos, S., Yim, S., Ryan, L. K., & Diamond, G. (2012). Induction of triggering receptor expressed on myeloid cells (TREM-1) in airway epithelial cells by 1,25(OH)2 vitamin D3. Innate Immunity, 18(2), 250–257. https://doi.org/10.1177/1753425911399796

Sassu, E. L., Kangethe, R. T., Settypalli, T. B. K., Chibssa, T. R., Cattoli, G., & Wijewardana, V. (2020). Development and evaluation of a real-time PCR panel for the detection of 20 immune markers in cattle and sheep. Veterinary Immunology and Immunopathology, 227, 110092. https://doi.org/10.1016/j.vetimm.2020.110092

Schauber, J., Dorschner, R. A., Coda, A. B., Büchau, A. S., Liu, P. T., Kiken, D., Helfrich, Y. R., Kang, S., Elalieh, H. Z., Steinmeyer, A., Zügel, U., Bikle, D. D., Modlin, R. L., & Gallo, R. L. (2007). Injury enhances TLR2 function and antimicrobial peptide expression through a vitamin D–dependent mechanism. Journal of Clinical Investigation, 117(3), 803–811. https://doi.org/10.1172/JCI30142

Scherberich, J. E., Kellermeyer, M., Ried, C., & Hartinger, A. (2005). 1-alpha-calcidol modulates major human monocyte antigens and toll-like receptors TLR 2 and TLR4 in vitro. European Journal of Medical Research, 10(4), 179–182.

Surlis, C., Earley, B., McGee, M., Keogh, K., Cormican, P., Blackshields, G., Tiernan, K., Dunn, A., Morrison, S., Arguello, A., & Waters, S. M. (2018). Blood immune transcriptome analysis of artificially fed dairy calves and naturally suckled beef calves from birth to 7 days of age. Scientific Reports, 8(1), Article 1. https://doi.org/10.1038/s41598-018-33627-0

Tabasi, N., Rastin, M., Mahmoudi, M., Ghoryani, M., Mirfeizi, Z., Rabe, S. Z. T., & Reihani, H. (2015). Influence of vitamin D on cell cycle, apoptosis, and some apoptosis related molecules in systemic lupus erythematosus. Iranian Journal of Basic Medical Sciences, 18(11), 1107–1111.

Takanashi, N., Tomosada, Y., Villena, J., Murata, K., Takahashi, T., Chiba, E., Tohno, M., Shimazu, T., Aso, H., Suda, Y., Ikegami, S., Itoh, H., Kawai, Y., Saito, T., Alvarez, S., & Kitazawa, H. (2013). Advanced application of bovine intestinal epithelial cell line for evaluating regulatory effect of lactobacilli against heat-killed enterotoxigenic Escherichia coli-mediated inflammation. BMC Microbiology, 13(1), 54. https://doi.org/10.1186/1471-2180-13-54

Tammaro, A., Derive, M., Gibot, S., Leemans, J. C., Florquin, S., & Dessing, M. C. (2017). TREM-1 and its potential ligands in non-infectious diseases: From biology to clinical perspectives. Pharmacology & Therapeutics, 177, 81–95. https://doi.org/10.1016/j.pharmthera.2017.02.043

Tomasinsig, L., De Conti, G., Skerlavaj, B., Piccinini, R., Mazzilli, M., D’Este, F., Tossi, A., & Zanetti, M. (2010). Broad-Spectrum Activity against Bacterial Mastitis Pathogens and Activation of Mammary Epithelial Cells Support a Protective Role of Neutrophil Cathelicidins in Bovine Mastitis. Infection and Immunity, 78(4), 1781–1788. https://doi.org/10.1128/IAI.01090-09

Villaggio, B., Soldano, S., & Cutolo, M. (2012). 1,25-dihydroxyvitamin D3 downregulates aromatase expression and inflammatory cytokines in human macrophages. Clinical and Experimental Rheumatology, 30(6), 934–938.

Wang, Q., He, Y., Shen, Y., Zhang, Q., Chen, D., Zuo, C., Qin, J., Wang, H., Wang, J., & Yu, Y. (2014). Vitamin D Inhibits COX-2 Expression and Inflammatory Response by Targeting Thioesterase Superfamily Member 4. Journal of Biological Chemistry, 289(17), 11681–11694. https://doi.org/10.1074/jbc.M113.517581

Wang, T.-T., Nestel, F. P., Bourdeau, V., Nagai, Y., Wang, Q., Liao, J., Tavera-Mendoza, L., Lin, R., Hanrahan, J. W., Mader, S., & White, J. H. (2004). Cutting Edge: 1,25-Dihydroxyvitamin D3 Is a Direct Inducer of Antimicrobial Peptide Gene Expression1. The Journal of Immunology, 173(5), 2909–2912. https://doi.org/10.4049/jimmunol.173.5.2909

Yamane, D., Kato, K., Tohya, Y., & Akashi, H. (2008). The relationship between the viral RNA level and upregulation of innate immunity in spleen of cattle persistently infected with bovine viral diarrhea virus. Veterinary Microbiology, 129(1), 69–79. https://doi.org/10.1016/j.vetmic.2007.11.004

Yang, Y., Bailey, J., Vacchio, M. S., Yarchoan, R., & Ashwell, J. D. (1995). Retinoic acid inhibition of ex vivo human immunodeficiency virus-associated apoptosis of peripheral blood cells. Proceedings of the National Academy of Sciences, 92(7), 3051–3055. https://doi.org/10.1073/pnas.92.7.3051

Zhang, Y., Leung, D. Y. M., Richers, B. N., Liu, Y., Remigio, L. K., Riches, D. W., & Goleva, E. (2012). Vitamin D Inhibits Monocyte/Macrophage Proinflammatory Cytokine Production by Targeting MAPK Phosphatase-1. The Journal of Immunology, 188(5), 2127–2135. https://doi.org/10.4049/jimmunol.1102412

